# Parallel social information processing circuits are differentially impacted in autism

**DOI:** 10.1101/2020.03.13.990549

**Authors:** Eastman M. Lewis, Genevieve L. Stein-O’Brien, Alejandra V. Patino, Romain Nardou, Cooper D. Grossman, Matthew Brown, Bidii Bangamwabo, Ndeye Ndiaye, Daniel Giovinazzo, Ian Dardani, Connie Jiang, Loyal A. Goff, Gül Dölen

## Abstract

Parallel processing circuits are thought to dramatically expand the network capabilities of the nervous system. Magnocellular and parvocellular oxytocin neurons have been proposed to subserve two parallel streams of social information processing, which allow a single molecule to encode a diverse array of ethologically distinct behaviors, although to date direct evidence to support this hypothesis is lacking. Here we provide the first comprehensive characterization of magnocellular and parvocellular oxytocin neurons, validated across anatomical, projection target, electrophysiological, and transcriptional criteria. We next used novel multiple feature selection tools in *Fmr1* KO mice to provide direct evidence that normal functioning of the parvocellular but not magnocellular oxytocin pathway is required for autism-relevant social reward behavior. Finally, we demonstrate that autism risk genes are uniquely enriched in parvocellular oxytocin neurons. Taken together these results provide the first evidence that oxytocin pathway specific pathogenic mechanisms account for social impairments across a broad range of autism etiologies.

**One Sentence Summary:** Pathoclisis of parvocellular oxytocin neurons plays an important role in the pathogenesis of social impairments in autism.

## Main Text

Parallel processing pathways have been identified in both sensory(1–6) and striatal(7– 9) circuits, and are thought to dramatically expand the network capabilities of the nervous system. Interestingly, oxytocin (OT) neurons are comprised of magnocellular and parvocellular subtypes (10) raising the possibility that this organizational principle can be extended to social information processing. Indeed, magnocellular and parvocellular OT neuronal subtypes can be differentiated by a number of characteristic features (11), which suggest that they are specialized to subserve distinct behaviors. For example, because magnocellular OT neurons are specialized to release large quantities of OT in both the central and peripheral nervous system, this adaptation may enable coordination of central and systemic responses during lactation and parturition (12). In contrast, parvocellular OT neurons release small quantities of OT restricted to the central brain, an adaptation that may subserve the measured reward associated with social cognition and peer-peer attachments (13). At this time this view is largely supported by indirect evidence from comparative studies across species (14). Nevertheless, the existence of parallel circuits for social information processing will have significant implications for understanding the pathogenesis of autism spectrum disorder (ASD), a neurodevelopmental disease characterized by domain specific impairments in social interactions (15).

We began using an established FluoroGold (FG) retrograde tracing approach (16) combined with oxytocin antibody labeling (17, 18) to carry out the first comprehensive quantification of the anatomical distribution and proportion of subtypes of OT neuron in the paraventricular nucleus (PVN) of the hypothalamus in male mice (postnatal day 30-40). OT-Ab labeled neurons were classified as magnocellular if they were FG positive, and parvocellular if they were FG negative (**Fig. S1A-F**). Quantification of these subtypes (n = 3 mice, every other section quantified) revealed that in the PVN, 66% (952 ± 69 neurons) of OTergic neurons are magnocellular, while 34% (488 ± 25 neurons) of OTergic neurons are parvocellular (**Fig. S1G**). Next, we examined the localization of all OT neurons, which revealed that magnocellular OT neurons are predominantly rostrally distributed compared to the caudally distributed parvocellular OT neurons (**Fig. S1H**).

Next, we sought to cross validate this finding by examining magnocellular and parvocellular electrophysiological signatures (19) in identified OT neurons using our recently generated OT-Flp recombinase driver line (20), crossed to a Flp-dependent GFP reporter line (21), (fdGFP) (**Fig. S2**). OT-2A-Flp::fdGFP mice (n = 9) were injected with i.v. FG, and GFP+ neurons were targeted for whole-cell current clamp recording in acute slices of the PVN (**Fig. 1A**). During recording, neurons were labeled with neurobiotin (Nb) to enable post-hoc classification of cells as FG positive (FG+, **Fig. 1B**) or FG negative (FG-, **Fig. 1C**). Neurons were depolarized from a hyperpolarized membrane potential (approximately −100 mV; **Fig. S3**) and the shape of the membrane depolarization preceding action potential (AP) initiation was classified as shoulder positive (Sh+, **Fig. 1D**; blue) or shoulder negative (Sh-, **Fig. 1D**; orange). OT neurons were examined for latency to first AP (**Fig. 1E**,**F**), AP duration (**Fig. 1H**,**I**), and sensitivity to the voltage-gated K+ antagonist 4-aminopyridine (4-AP, **Fig. 1J-O**). These studies confirm that latency to first AP is longer in Sh+ compared to Sh-OT neurons (**Fig. 1E**), as well in FG+ compared to FG-OT neurons when neurons are grouped by FG labeling (**Fig. 1F**). Furthermore, AP duration (**Fig. 1G**) is longer in Sh+ compared to Sh-OT neurons (**Fig, 1H**) as well as longer in FG+ versus FG-OT neurons (**Fig. 1I**). Application of the voltage-gated K+ channel blocker 4-AP (5 mM; **Fig. 1J**,**M**) reduced the latency to first AP and increased AP duration in magnocellular OT neurons, but had no effect on latency in parvocellular OT neurons (**Fig. 1K,L,N,O**).

**Fig. 1.**
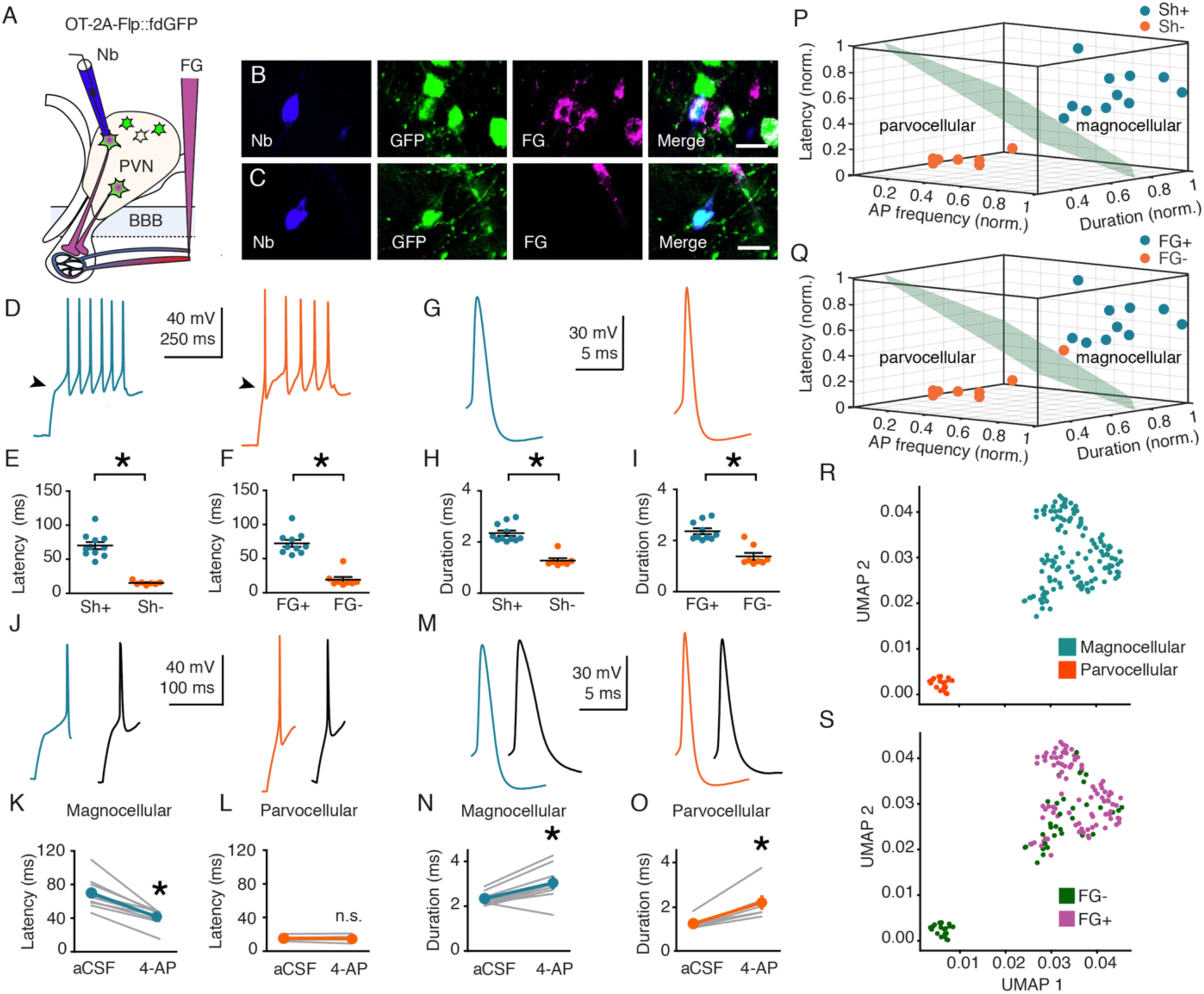
Electrophysiological signatures of magnocellular and parvocellular OT neurons. **A**, Validation strategy using retrograde labeling in the tail vein and Nb-filled recording pipette. **B**,**C**, FG+ (B) and FG- (C) neurons recovered after electrophysiological recording. Nb (blue, left), GFP (green, center-left), FG (magenta center-right), merge (right). Scale bar 20 µm. **D**,**G**, Representative traces from magnocellular (blue), and parvocellular (orange) OT neurons. **E**,**F**,**H**,**I**, Latency to first AP and AP duration are shorter in Sh-compared to Sh+ OT neurons (t_(16)_ = 8.286; p > 0.0001 (latency), t_(16)_ = 6.987; p > 0.0001 (duration)) (E,H) and FG-compared to FG+ OT neurons (t_(16)_ = 7.941; p < 0.0001 (latency), t_(16)_ = 5.524 p > 0.0001 (duration)) (F,I). **J**,**M**, Representative traces (blue, magnocellular OT; orange, parvocellular OT; black, following 5 mM 4-AP). **K**,**L**, 4-AP reduces latency to first AP in magnocellular (t_(9)_ = 6.678; p < 0.0001), (K) but not parvocellular (t_(6)_ = 0.998; p = 0.357) (L) OT neurons. **N**,**O**, 4-AP increases AP duration in magnocellular (t_(9)_ = 4.491; p = 0.002) (N) and parvocellular (t_(6)_ = 5.017; p = 0.002) (O) OT neurons. **P**,**Q**, K-means cluster analysis using z-scored values for AP frequency, duration, and latency to first AP for each neuron. **P**, Clusters coincided completely (100%, 18 of 18 neurons) with qualitative magnocellular or parvocellular categorization based on shoulder. **Q**, Cluster assignment matched magnocellular or parvocellular identity of 94% (17 of 18) neurons defined by FG labeling. **R**, UMAP of scRNASeq data delineates two clusters indicating two distinct neuron types, magnocellular (blue) and parvocellular (orange). **S**, Distribution of FG labeling confirms putative cell type designations. The parvocellular cluster is devoid of fluorogold positive cells, (21 of 21 cells; 100% FG-) and the magnocellular cluster is majority FG+ (91 of 125 cells; 72.8% FG+). Note that within the magnocellular cluster, FG negative cells display markers of low RNA quality (**Fig. S4**). Data represented as mean ± sem. * indicates p < 0.05; Student’s t-test (two-tailed; paired (K,L,N,O) two-tailed; unpaired (E,F,H,I)) for statistical comparisons.

To quantify whether an individual OT neuron’s magnocellular or parvocellular identity can be predicted from its electrophysiological features, we used unsupervised k-means clustering analysis. The resulting clusters corresponded completely (100%, 18 of 18 OT neurons) with the qualitative magnocellular or parvocellular categorization based on the presence or absence of a shoulder in the electrophysiological trace (**Fig. 1P**) and in 94% (17 of 18 OT neurons) when defined by FG labeling, (**Fig. 1Q**). The proportions of magnocellular and parvocellular neurons obtained using these methods are consistent (**Fig. S3**) with those determined in **Fig. S1**. Taken together, these studies demonstrate that magnocellular and parvocellular subtypes of OT neurons can be reproducibly defined by cross-validated electrophysiological and anatomical criteria.

In order to determine whether magnocellular and parvocellular OT neurons exhibit distinct transcriptional profiles, next we employed a single cell RNA-Seq approach in OT-2A-Flp::fdGFP mice injected with i.v. FG as described above (**Fig. S4**). In total 194 OT neurons were collected by fluorescence assisted cell sorting (FACS) and used as input for a modified SMART-Seq2 scRNA-Seq library preparation as described previously (22). Using density peak clustering, we identified two transcriptionally distinct clusters of neurons (**Fig. 1R**). To annotate clusters as magnocellular or parvocellular OT neurons, we assessed the distribution of FG labeling across the two clusters (**Fig. 1S**). The smaller cluster (21 neurons) consisted entirely of FG-cells across all three replicates (**Fig. S4**), consistent with a parvocellular identity. Conversely, the majority of cells (72.8%) in the larger cluster were FG+ across replicates (**Fig. S4**), consistent with a magnocellular identity. While our electrophysiological and anatomical cross validation suggests that nearly all magnocellular OT neurons should be labeled by i.v. FG (**Fig 1Q**), a positive correlation was observed between FG labeling and cellular quality metrics, suggesting that our ability to detect FG labeling may be negatively affected by the cellular dissociation. Consistent with this interpretation, we observed that parvocellular OT neurons displayed significantly reduced intra-cell type heterogeneity relative to magnocellular OT neurons (**Fig. S4**).

Using the monocle2 likelihood ratio test (23), we identified 181 genes (**Table S1**) with significant (0.1% FDR; Monocle2; BH-corrected) differential expression (DE) between magnocellular and parvocellular OT neurons (**Fig. 2A**), representing novel discriminating marker genes for these two subtypes. Amongst the most significantly differentially expressed genes was the calcium-binding protein Calbindin (*Calb1*) and a large conductance calcium-activated potassium channel subunit (*Kcnmb4*), both enriched for expression in the magnocellular OT neuron population (**Fig. 2B)**, as well as the extracellular matrix serine protease Reelin (*Reln*) and the cannabinoid receptor 1 (*Cnr1*) genes, both uniquely expressed in parvocellular OT neurons (**Fig. 2C**). These results were validated using anatomical criteria (**Fig. S1**) combined with fluorescent *in situ* hybridization chain reaction version 3.0 (24) (HCR 3.0; **Table S2**), which demonstrated that *Oxt+* neurons in the rostral portion of the PVN were specifically colocalized with *Calb1* and not *Cnr1* (**Figure 2D**), while in more caudal sections *Oxt+* neurons colocalized with *Cnr1* and not *Calb1* (**Figure 2E**). These results are the first to establish a transcriptional signature for magnocellular and parvocellular OT neuronal subtypes.

**Fig. 2.**
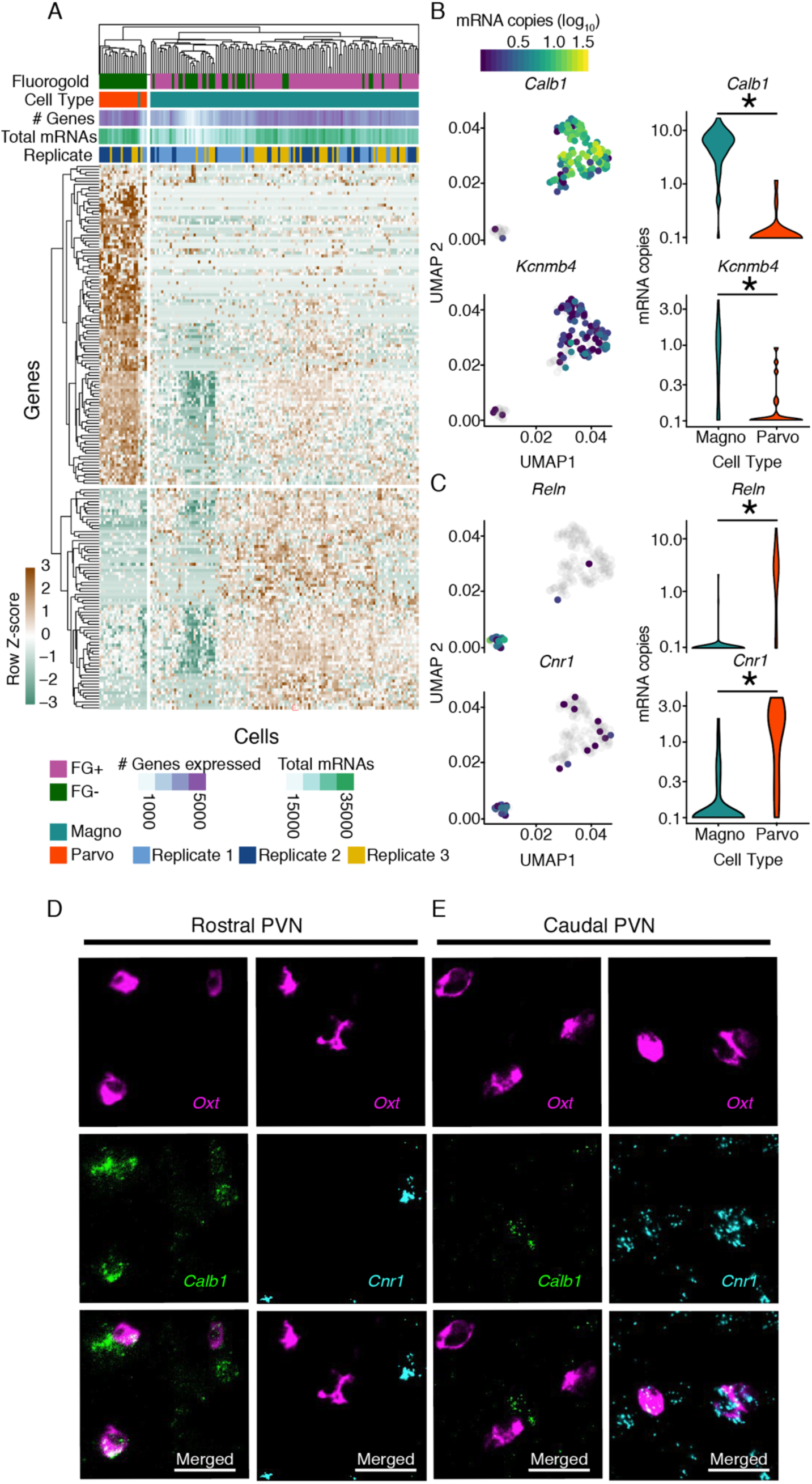
Differential expression analysis of clusters determines unique molecular markers of magnocellular and parvocellular OT neurons. **A**, Heatmap of significantly differentially expressed genes (Monocle Likelihood Ratio Test, FDR 0.1%) between magnocellular and parvocellular OT neurons. **B**, UMAPs (left) and Violin plots (right) for genes expressed more highly in magnocellular OT neurons: *Calb1* and *Kcnmb4* (identified using FDR 0.00001%; p = 1.28e-^30^ and p = 5.46e-^7^, respectively). **C**, UMAPs (left) and Violin plots (right) for genes expressed more highly in parvocellular OT neurons: *Reln* and *Cnr1* (identified using FDR 0.00001%; p = 1.48e-15 and p = 6.67e-^11^, respectively). **D**,**E**, Validation of differential expression via fluorescent *in situ* hybridization chain reaction version 3.0 (HCR 3.0) within the PVN. *Calb1* but not *Cnr1*, is colocalized with *Oxt* within the rostral PVN (D). *Cnr1*, but not *Calb1* is colocalized with *Oxt* in the caudal PVN (E). Scale bars 20 µm. Distribution of *Calb1* (rostral) and *Cnr1* (caudal) expression in OT neurons recapitulates differential localization of magnocellular (rostrally enriched) and parvocellular (caudally enriched) neurons respectively. * indicates p < 0.05; Monocle2 likelihood ratio test for statistical comparisons.

We next sought to test the hypothesis that these neuronal subtypes subserve discrete functions that may be differentially impacted in the Fragile X subtype of autism. We began by characterizing three forms of reward learning in the constitutive *Fmr1* knockout (KO) (25) using peer-peer social, alloparent social, and cocaine conditioned place preference (CPP) assays (26–28). These results demonstrate a significant impairment in peer-peer social reward learning in *Fmr1* KO mice compared to wildtype (WT), while both WT and *Fmr1* KO mice exhibit significant cocaine CPP **(Fig. S5E-H)**, as well as alloparent CPP **(Fig. S5I-L)**. Importantly, the social domain specificity of this phenotype in the *Fmr1* KO ASD model mouse is consistent with the social motivation theory of autism, which posits that characteristic social impairments, including failure to develop peer relationships, difficulty with emotional reciprocity and imitative play, and impaired cognitive empathy are the consequence of a domain specific derailment in peer-peer social reward learning (15). Furthermore, this is the pattern of reward learning deficits that has been hypothesized a selective impairment in parvocellular OT signaling would produce (14).

Additionally, this impairment in peer-peer social CPP in the *Fmr1* KO mouse recapitulates previously reported consequences of OT receptor blockade or ablation in the Nucleus Accumbens (NAc) (29), raising the possibility that these phenotypes share overlapping mechanisms. In order to test whether magnocellular or parvocellular OT neurons subserve this function, we next injected the NAc of OT-2A-Flp::fdGFP mice (n = 6 mice) with retrogradely transported fluorescent microspheres (retrobeads; RtB (30)), and performed current clamp recordings from RtB/GFP double-positive neurons in acute slices of the PVN (**Fig. 3A**,**B**). All Rtb labeled, OT positive neurons recorded (9 of 9) exhibited a parvocellular electrophysiological phenotype by latency to first AP, AP duration, 4-AP sensitivity, and K-means clustering signatures (**Fig. 3C-K**). Taken together with the impairment in peer-peer social reward learning seen in *Fmr1* KO mice, these findings are consistent with the hypothesis that parvocellular OT neurons subserve a discrete and parallel set of social behaviors (14).

**Fig. 3.**
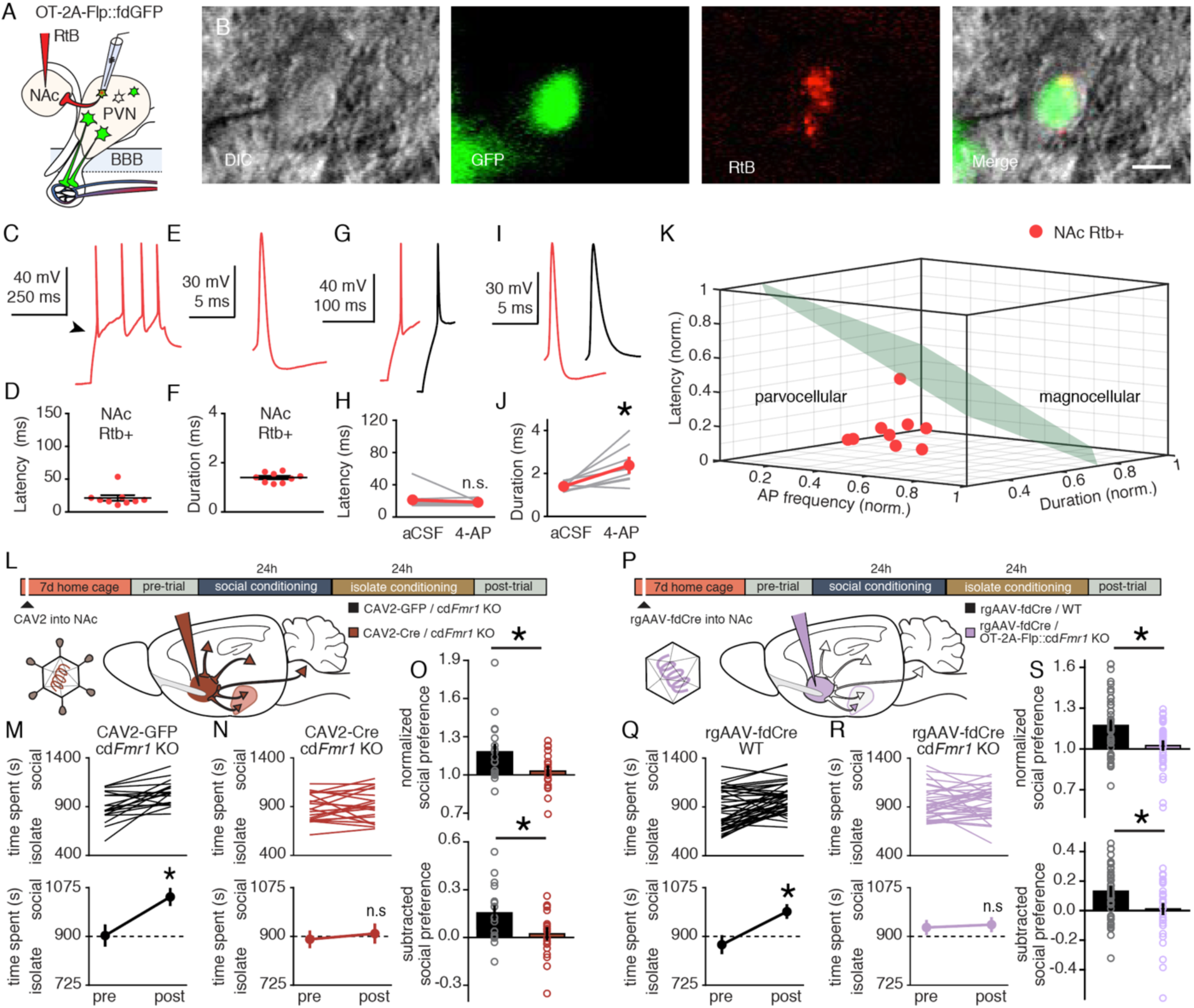
Parvocellular OT neurons innervate the NAc and require *Fmr1* to mediate social reward learning. **A**, RtB labeling and recording strategy in OT-2A-Flp::fdGFP mice. **B**, Example of a GFP/RtB double-positive OT neuron targeted for electrophysiological recording. Left to right; DIC (gray), RtB (red), GFP (green), merge. Scale bar 10 µm. **C**,**E**, Traces recorded from an NAc-projecting OT neuron. **D**, Latency to first AP of NAc-projecting OT neurons (n = 9; 21.2 ± 4.2 ms). **F**, AP duration of NAc-projecting OT neurons (n = 9; 1.4 ± 0.07 ms). **G**,**I**, Traces (red, RTB+; black, following 5 mM 4-AP). **H**,**J**, 4-AP does not change latency to first AP (t_(7)_ = 0.981; p = 0.359) (H), but increases AP duration (t_(7)_ = 3.193 p = 0.015) (j) in NAc-projecting OT neurons. **K**, Using the k-means clustering algorithm trained using data from Fig. 1, 100% (9 of 9) sampled RtB/GFP double-positive cells were assigned to the parvocellular cluster. **L**, Protocol for social CPP following acute knockdown of *Fmr1* in neurons projecting to NAc. **M**,**N**, Individual (top) and average (bottom) data for cd*Fmr1* KO mice injected with CAV2-GFP (M) and CAV2-Cre (N). cd*Fmr1* KO mice injected with CAV2-GFP (t_(18)_ = 3.95; p = 0.001) but not CAV2-Cre (t_(19)_ = 0.69; p = 0.499) exhibit social CPP. **O**, Normalized (t_(37)_ = 2.453; p = 0.019) and subtracted (t_(37)_ = 2.566; p = 0.015) comparisons reveal reduced magnitude social CPP in cd*Fmr1* KO mice injected with CAV2-Cre compared to CAV2-GFP. **P**, Protocol for social CPP following acute knockdown of *Fmr1* in OT neurons projecting to NAc. **Q**,**R**, Individual (top) and average (bottom) data for WT (Q) and OT-2A-Flp::cd*Fmr1* KO (R) mice injected with rgAAV-fdCre. WT (t_(43)_ = 4.947; p > 0.0001) but not OT-2A-Flp::cd*Fmr1* KO (t_(35)_ = 0.349; p = 0.729) mice injected with rgAAV-fdCre exhibit social CPP. **S**, Normalized (t_(78)_ = 3.295; p = 0.002) and subtracted (t_(78)_ = 2.937; p = 0.004) comparisons reveal reduced magnitude social CPP in OT-2A-Flp::cd*Fmr1* KO compared to WT mice injected with rgAAV-fdCre. Data represented as mean ± sem. * indicates p < 0.05; Student’s t-test (two-tailed; paired (H,J,M,N,Q,R) two-tailed; unpaired (O,S)) for statistical comparisons.

In order to directly test the behavioral consequence of disrupting the parvocellular OT input to the NAc, we began by testing whether ongoing *Fmr1* function is required for peer-peer social reward learning. We utilized conditional (Cre dependent) *Fmr1* KO mice (31) (cd*Fmr1* KO) receiving injections of Canine Adenovirus expressing Cre (CAV-Cre) or GFP (CAV-GFP) (32, 33) bilaterally into the NAc. Seven days later, social CPP was measured and revealed impaired social CPP in cd*Fmr1* KO mice injected with CAV-Cre, but not CAV-GFP (**Fig. 3L-O**). To test the OT cell type specific role of FMRP (the protein encoded by the *Fmr1* gene) in regulating social reward learning, we crossed cd*Fmr1* KO mice to OT-2A-Flp mice, and generated retrograde Adeno-Associated Virus expressing Flp-dependent-Cre (rgAAV-fdCre)(34–36) (**Fig. 3P**). Consistent with a required and ongoing role for FMRP in OT neurons projecting to the NAc specifically, social CPP was absent in OT-2A-Flp::cd*Fmr1* KO, while WT animals injected with the same virus exhibited normal social reward learning (**Fig. 3Q-S**). Injection sites and viral function were confirmed as shown in **Fig. S6**. To exclude the possibility that knockdown of *Fmr1* in magnocellular OT neurons would produce the same effect, we capitalized on our discovery of *Calb1* as a marker of magnocellular OT neurons and crossed *Calb1*-IRES-Cre mice with cd*Fmr1* KO mice to selectively knock down *Fmr1* in neurons expressing *Calb1*. Consistent with the unique requirement of *Fmr1* in parvocellular OT neurons for peer-peer social reward learning, social CPP was not impaired in *Calb1*-IRES-Cre::cd*Fmr1* KO mice (**Fig. S7**). Taken together, these anatomical, electrophysiological, and functional genomic data demonstrate that social reward learning deficits seen in constitutive *Fmr1* KO mice are not the consequence of hysteresis, but rather the ongoing requirement of *Fmr1* in parvocellular, but not magnocellular OT neurons.

Having identified a role for *Fmr1* in parvocellular OT neuronal regulation of domain specific social reward learning, we next sought to determine whether this pathogenic mechanism in Fragile X may generalize to other ASD etiologies. To do this, we performed gene set enrichment analysis for ASD risk genes (37) and FMRP binding partners (38). In both cases, genes were significantly enriched in parvocellular relative to magnocellular neurons (**Fig. 4A,B**; **Tables** S**3 and S4**). Additionally, 10 of the 12 differentially expressed genes in the intersection of the 2 groups (*Dlgap2, Cnr1, Pacs1, Tanc2, Anks1b, Slc6a1, Synj1, Tspan7, Camk2b, Stxbp1, Fbxo11, Foxp1*; **Fig. 4A,B**, individually labeled genes; **Fig. S8**) were also enriched in parvocellular cells. Since OT neurons in both the rostral and caudal PVN (**Fig. S8**) express FMRP, these results indicate that subtype specific expression of FMRP itself is unlikely to account for selective enrichment of ASD risk genes and FMRP binding partners in parvocellular OT neurons. Next we repeated this analysis on risk genes for Schizophrenia, Alzheimer’s Disease, and Type II Diabetes (39–41). While a handful of Schizophrenia risk genes were differentially expressed in OT neuronal subtypes, these were not significantly enriched in either magnocellular or parvocellular OT neurons (p = .98, **Fig. 4C**). Moreover, Alzheimer’s Disease had only a single intersecting gene and Type II Diabetes risk genes were completely absent amongst differentially expressed genes in OT cells (**Fig. S9**). These results indicate that the cell type specific vulnerability of parvocellular OT neurons to genetic injury generalizes across multiple ASD etiologies, but is not apparent in unrelated genetic disorders that are not defined by impairments in social behavior.

**Fig. 4.**
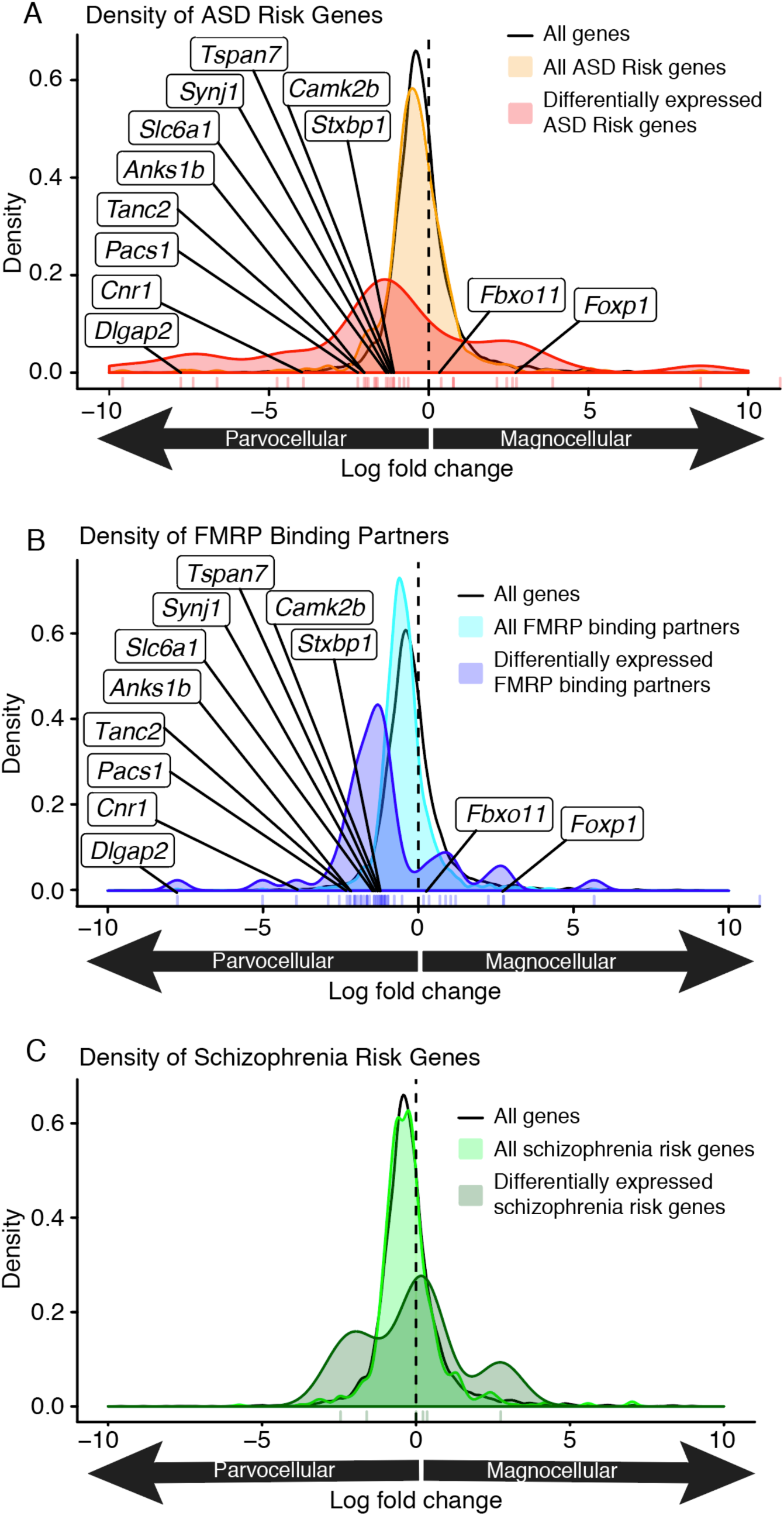
ASD risk and FMRP target genes are enriched in parvocellular OT neurons. **A**, Density of the log2 fold change for (black) all expressed genes, (orange) all ASD risk genes, and (red) all significantly differentially expressed (FDR = 0.1%) ASD risk genes. ASD risk genes (orange vs, black; p = 0.002041; one-sided Wilcoxon Rank Sum Test) and differentially expressed ASD risk genes (red vs black; p = 0.0002611) are enriched in parvocellular OT neurons. Individually labeled genes are the union of ASD risk genes and FMRP binding partners that are differentially expressed between neuron types. **B**, Density of the log2 fold change for (black) all expressed genes, (light blue) all FMRP binding partners, and (dark blue) all significantly differentially expressed (FDR 0.1%) FMRP binding partners. FMRP binding partners (light blue vs black; p = 6.83e^-16^; one-sided Wilcoxon Rank Sum Test) and differentially expressed FMRP binding partners (dark blue vs black; p = 1.751e^-9^) are enriched in parvocellular OT neurons. Individually labeled genes are the union of ASD risk genes and FMRP binding partners that are differentially expressed between neuron types (FDR 0.1%). **C**, Density of the log2 fold change for (black) all expressed genes, (light green) all schizophrenia risk genes, and (dark green) all significantly differentially expressed (FDR 0.1%) schizophrenia risk genes. Schizophrenia risk genes are not enriched in magnocellular or parvocellular OT neurons (light green vs black; p = 0.98) and differentially expressed schizophrenia risk genes (dark green vs black; p = 0.21; one-sided Wilcoxon Rank Sum Test) are not enriched in parvocellular OT neurons. Vertical tick marks below density plots indicate the individual differentially expressed ASD risk genes (A), FMRP binding partners (B), and schizophrenia rick genes (C).

In recent years intranasal OT administration, which likely targets magnocellular signaling pathways (42), has been employed widely in clinical trials in an attempt to ameliorate social behavioral deficits in ASD. To date, these clinical trials remain inconclusive, perhaps due to the limited understanding of the parallel actions of magnocellular and parvocellular OT neurons (14). Here we provide evidence that parvocellular OT neurons are the source of OTergic innervation of NAc that supports autism relevant social reward learning, and that parvocellular, rather than magnocellular OT neurons are vulnerable to ASD-relevant genetic insults (29). Together these results emphasize functional importance of distinct OT neuron subtypes, and suggest that strategies that target parvocellular OT signaling mechanisms may serve as a more appropriate and effective therapeutic intervention than peripheral OT administration. Beyond a specific role of OT in the pathogenesis of social deficits in ASD, our results highlight the importance of using circuit-level transcriptional analysis to understand the cause of behavioral impairments in complex genetic disorders. Many disorders of brain function contain an element of genetic heritability (43), which can be difficult to reconcile with highly specific behavioral deficits. Building upon the discovery that unique molecular features can confer vulnerability to neurons within a specific brain nucleus to genetic lesion (44), our results demonstrate this same mechanism of pathoclisis can impact discreet circuit elements comprised of specific neuronal subpopulations within a given nucleus. Rather than excluding the possibility that similar mechanisms elsewhere in the brain contribute to behavioral deficits in psychiatric illness, we propose that our findings here outline a framework for reconciling the consequences broad genetic insults with discreet behavioral deficits that arise from circuit-level neuronal dysfunction.

## Acknowledgments

We thank members of the Dölen and Goff labs for comments on the manuscript, Elizabeth Vincent for histological consultation, Ben Emmert and Arjun Raj (UPenn) for assistance with the HCR 3.0 protocol. PS38 and VA10 antibodies were kind gifts from Dr. Harold Gainer.

## Author contributions

E.M.L., A.V.P., G.L.S.O., L.A.G., and G.D. designed the experiments and wrote the manuscript; E.M.L. and C.D.G. conducted electrophysiology and analyzed the data. E.M.L, R.N, A.V.P and N.N. conducted behavioral experiments and analyzed the data. E.M.L., A.V.P. and B.B. conducted immunoblotting experiments and analyzed the data. A.V.P. and G.L.S.O. conducted the scRNAseq experiments. G.L.S.O. and L.A.G. analyzed the scRNAseq data. A.V.P. and G.L.S.O performed the HCR smFISH under the direction of I.D. and C.J. I.D. performed smFISH imaging.

## Data and materials availability

OT-2A-Flp mice will be made available with a materials transfer agreement. The data and code that support the findings of this will be made available upon request to the corresponding authors. Genetic data will be publicly available via the NCBI Gene Expression Omnibus upon publication of the manuscript in accordance with CZI guidelines.

## Supplementary Materials

Materials and Methods

Figures S1-S10

Tables S1-S4

References (45-53)

