## supplementary material for "Parallel social information processing circuits are differentially impacted in autism"

**This PDF file includes:**

Materials and Methods
Figs. S1 to S10
Tables S1 to S4

**Materials and Methods**

Animals

Experiments were conducted in 3 - 7 week-old male mice. Wild-type C57BL6/J (Stock # 000664; 3 - 4 weeks old) and *Calb1*-IRES-Cre (Stock # 028532) mice were obtained from Jackson Laboratories. Flp-dependent GFP reporter mice (Stock # 32038) were obtained from the Mutant Mouse Resource Center. Transgenic mice were bred in-house and weaned at 3 weeks of age. *Oxytocin*-2A-Flp-optimized (OT-2A-Flp) knockin mice were generated by Cyagen Biosciences (Santa Clara, CA) (**Supplementary Fig. 2**)(*20*). Similar to the strategy employed by the oxytocin-IRES-Cre knockin mouse (*45*) the stop codon in exon 3 of the endogenous OT gene was replaced with a 2A-Flp-optimized (FlpO) construct. The co-translational cleavage 2A peptide strategy (known to have a more robust expression of its downstream gene compared to IRES (*46*)) has been used to separate Flp and OT, since Flp is directed to the nucleus, while OT is directed to the cytoplasm. OT-2A-Flp mice were kept on a C57Bl/6 background. To generate hemizygous *Fmr1* KO males for behavior, homozygous *Fmr1* KO females were bred to hemizygous *Fmr1* KO males. The same strategy was used to generate Cre-dependent conditional *Fmr1* KO (cd*Fmr1* KO) males. To generate OT-2A-Flp::Ai9 and OT-2A-Flp::cd*Fmr1* KO mice, homozygous OT-2A-Flp males were crossed with homozygous Ai9 or cd*Fmr1* KO females respectively. To generate *Calb1*-Cre::cd*Fmr1* KO males for behavior, homozygous *Calb*1-IRES-Cre males were crossed with homozygous cd*Fmr1* KO females. Homozygous or heterozygous OT-2A-Flp and fdGFP mice were crossed to generate OT-2A-Flp::fdGFP mice. All mice were maintained on a on a 12:12h natural light dark cycle, starting at 7:30am with food and water ad libitum. All procedures were conducted in accordance with protocols approved by the Johns Hopkins Animal Care and Use Committee

Genotyping

OT-2A-Flp knockin mice, fdGFP, *Fmr1* KO, and cd*Fmr1* KO mice were genotyped using polymerase chain reaction (PCR) analysis of DNA isolated from tail snips taken before weaning. For OT-2A-Flp mice, the wild-type allele was identified by a 622-bp PCR product and the mutant allele by a 308-bp PCR product using the primers mOxtR1: TCCGACAATTAGACACCAGTCAA; mOxtF1: CTACCTGAGCAGCTACATCAACAG; mOxtF2: AGGGCTTTGGGAAGTGTTAGGCT. The reactions were run under the following conditions: 94°C × 3 min, (94°C × 30 s, 60°C × 35 s, 72°C × 35 s) × 38 cycles, 72°C × 5 min. For fdGFP mice, the wild-type allele was identified by a 603-bp PCR product, and the mutant allele by a 320-bp PCR product using the following primers; mutant: CCA GGC GGG CCA TTT ACC GTA AG; common: AAA GTC GCT CTG AGT TGT TAT; wild-type: GGA GCG GGA GAA ATG GAT ATG. The reactions were run using the Gt(ROSA)26Sor$^{tm(CAG)}$ protocol published by Jackson Laboratories. For the *Fmr1* KO mice, the wild-type allele was identified by a 131-bp PCR product, and the mutant allele by a 400-bp PCR product using the following primers; wild-type: TGT GAT AGA ATA TGC AGC ATG TGA; common: CTT CTG GCA CCT CCA GCT T; mutant: CAC GAG ACT AGT GAG ACG TG. The reactions were run using the Fmr1$^{tm1Cgr}$ protocol published by Jackson Laboratories. For cd*Fmr1* KO mice, the wild-type allele was identified by a 120-bp PCR product, and the mutant allele was detected by a 220-bp PCR product using the following primers: CCC ACT GGG AGA GGA TTA TTT GGG and GTT GAG CGG CCG AGT TTG TGA G. Reactions were run under the following conditions: 95°C × 3 min, (95°C × 30 s, 55°C × 30 s, 72°C × 60 s) × 35 cycles, 72°C × 7 min.

Social Conditioned Place Preference

The social CPP assay was conducted as previously described(*20*). Briefly, animals were weaned and socially housed (3–5 same sex cage mates; **Fig. S10**) in a cage containing corncob bedding (Anderson Cob, 1/4" cob or 1/8" cob; Animal Specialties and Provisions). At approximately 6 weeks of age, animals were placed in an open field activity chamber (ENV-510, Med Associates) employing infrared beams and a software interface (Activity Monitor, Med Associates) to monitor the position of the mouse. The chamber was partitioned into two equally sized zones using a clear Plexiglas wall; each zone containing one type of novel bedding (Alpha-Dri; Animal Specialties and Provisions, Kaytee Soft Granule; Petco, Anderson Cob, 1/4" cob; Animal Specialties and Provisions, or Aspen Chip; Northeastern Products). To establish each animal's baseline preference for the bedding cues, the amount of time each mouse spent exploring each zone was recorded during a 30 min pre-conditioning trial. Immediately after this trial, mice received social conditioning with cage mates for 24 hr on one type of bedding, followed by 24 hr isolation conditioning on the other bedding. Immediately following the isolation conditioning, a 30 min post-conditioning trial was conducted to establish preference for the two conditioned cues. Chamber assignments were counterbalanced for side and bedding cues. Exclusion criteria for this behavior were defined as a pre-conditioning preference score $\left(time\ side\ 1/\frac{total\ time}{2}\right)$ of > 1.5 or < 0.5. Experimental conditions were compared using normalized social preference scores $(\frac{social\ zone\ post}{social\ zone\ pre})$, and subtracted social preference scores $(\frac{(social\ zone\ post - social\ zone\ pre)}{900})$ (*20*, *29*).

Cocaine Conditioned Place Preference

The protocol for cocaine CPP was conducted as previously described (*20*). Experiments were performed in the same open field activity chambers as social CPP using an identical configuration.

After 3 days of habituation to i.p. saline injections in the home cage, the pre-conditioning trial was performed as stated above. After 24 h, mice received an i.p. injection of cocaine (20 mg/kg) immediately followed by 30 min conditioning on one bedding (Soft Granule or Alpha Dri), which was randomly assigned in a counterbalanced fashion. A second 30 min conditioning session was conducted 24 h later on the other bedding after an i.p. injection of saline (equal volume to cocaine). A 30 min post-conditioning test session was conducted 24 h later to determine each mouse's preference for the cocaine versus saline associated beddings.

<u>Alloparent Conditioned Place Preference</u>

The alloparent CPP assay was adapted from a previously used assay (*28*), and experiments were conducted in the same open field activity chambers using an identical configuration to the social CPP assay. To generate familiar alloparent stimulus animals, adult (> 9weeks old) virgin wild type C57BL6/J females were pair housed with pregnat WT or *Fmr1* KO females. This housing configuration remained intact following the birth of WT or *Fmr1* KO pups until alloparernt CPP testing began. Alloparent conditioning was conducted as follows. Beginning at postnatal day 19-20, and continuing until each male littermate was tested, male mice were individually weaned from their home cage and placed directly in an activity chamber with two novel beddings. To establish each animal's baseline preference for the novel bedding cues, the amount spent exploring each zone of the activity chamber was recorded during a 30 min pre-conditioning trial. Immediately after this trial, mice (postnatal day 19-24) individually received alloparent conditioning with their virgin female alloparent for 24 hr on one type of bedding, followed by 24 hr isolation conditioning on the other bedding. Immediately following the isolation conditioning, a 30 min post-conditioning trial was conducted to establish preference for the two conditioned cues. This process was repeated individually until each male littermate had been tested. Identical to the methods used for social CPP, chamber assignments were counterbalanced for side and bedding cues. Exclusion criteria for this behavior were defined as a pre-conditioning preference score $\left(time\ side\ 1/\frac{total\ time}{2}\right)$ of > 1.5 or < 0.5. Experimental conditions were compared using normalized social preference scores ($\frac{social\ zone\ post}{social\ zone\ pre}$), and subtracted social preference scores ($\frac{(social\ zone\ post - social\ zone\ pre)}{900}$) (*20*, *29*).

Stereotaxic and tail-vein injections

All stereotaxic injections into the nucleus accumbens (NAc; distance from bregma: anterior +1.54 mm; lateral $\pm$1.065 mm; ventral - 4.1 mm)) were performed under general ketamine-medetomidine anesthesia using a stereotaxic instrument (David Kopf). For electrophysiological analysis of NAc-projecting OT neurons, a small volume (~30 nl) of diluted Rtb solution (1:4; Lumafluor; Red Retrobeads) was injected unilaterally or bilaterally into NAc core at a slow rate (20 nl/min) using a syringe pump (Harvard Apparatus, MA). For conditional knockdown of *Fmr1*, 1 µl viral suspension (0.1 µl/min; CAV2-GFP, CAV2-Cre-GFP, rgAAV-fDIO-Cre-GFP, or rgAAV-fDIO-Cre-HA) was injected bilaterally into NAc 7 days before behavioral testing. CAV2-GFP and CAV2-Cre-GFP viruses were obtiained from Plateforme de Vectorologie de Montpellier. The fDIO-Cre-GFP plasmid was a gift of Dr. Bo Li and Dr. Linda Van Aelst and viral packaging was conducted by Stanford University's Neuroscience Gene Vector and Virus Core. rgAAV-fDIO-Cre-HA was obtained from Addgene (Catalog # 121675-AAVrg). To confirm viral-mediated Cre function, 1 µl CAV2-Cre-GFP or 1.5 µl rgAAV-fDIO-Cre-HA was injected into NAc of Ai9 (tdTomato Cre-reporter) or OT-2A-Flp::Ai9 mice respectively. After all injections, the injection pipette was left in place for at least 5 min prior to removal from the brain. Injection sites were confirmed post-hoc by preparing sections (20-50 µm) containing the NAc. For i.v.

labeling of magnocellular OT neurons with FG, 1 or 2 tail-vein injections of 15 µl 4% FG (Fluorochrome) diluted in sterile saline were performed using a syringe and 25-gauge needle at least 24 hours prior to sacrifice. To facilitate i.v. injections, mice were placed under a warm lamp for 3-5 min and were briefly restrained using a tail vein restrainer (TV-150; Braintree Scientific).

Electrophysiology

Using a Leica VT 1200s vibrating microtome, coronal sections (250 µm) containing the paraventricular nucleus were cut in an ice-cold sucrose solution containing the following (in mM); Sucrose 228; $NaHCO_3$ 26; Glucose 11; KCl 2.5; $NaH_2PO_4$ 1; $MgSO_4$ 7; $CaCl_2$ 0.5. Immediately after cutting, slices were transferred into 32˚C aCSF containing (in mM); NaCl 119; KCl 2.5; $NaH_2PO_4$ 1; $NaHCO_3$ 26.2; Glucose 11; $MgCl_2$ 1.3; $CaCl_2$ 2.5. Slices were allowed to recover at 29˚C for at least 1 hour prior to recording. GFP+ and RtB+ neurons were visualized using 470 nm and 535 nm LEDs respectively (Cool LED). LED excitation was delivered through a 40x microscope objective, (Olympus) and fluorescence was detected using a camera and visualized using SliceScope Pro software (Scientifica). Recordings were conducted in current clamp using patch pipettes (2-4 MΩ) filled with internal solution containing the following (in mM: 130 K-Gluconate, 10 HEPES, 1 NaCl, 1 $CaCl_2$, 10 EGTA, 1 MgCl, 2 Mg-ATP, 0.5 Na-GTP, and 0.125 % Neurobiotin) and the liquid junction potential (-13 mV) was corrected after recording. Recording was discontinued for neurons that failed to maintain a hyperpolarized resting membrane potential in the absence of current injection, and neurons that failed to produce APs > 50 mV were excluded from further analysis. Data was acquired and analyzed using the Recording Artist plugin in Igor Pro and custom software written in MATLAB (Mathworks). AP initiation was determined as the first point where the membrane potential accelerated past 40 $ms/s^2$. This threshold was confirmed with comparison to manual determination of AP initiation. AP peaks were determined by finding local maxima with a minimum peak of 0 mV and a minimum separation of 5 ms. Nine mice, with a maximum of 4 neurons per animal were used to cross-validate FG and electrophysiological differentiation of magnocellular and parvocellular neurons. Six mice, with a maximum of 2 neurons per animal were used for electrophysiological characterization of OTergic projections to NAc.

Cell Isolation, Enrichment, and cDNA Library Preparation

OT-2A-Flp::fdGFP mice were weaned at P21 and given i.v. injections of FluoroGold on that and the following day. At P23 mice were sacrificed. Brains were rapidly removed and 250 µm thick coronal slices (n = 7 mice) containing the paraventricular nucleus of the hypothalamus were sectioned using a Leica VT-1200s vibrating microtome in ice-cold ACSF solution containing the following (in mM): 124 NaCl; 2.5 KCl; 1.2 NaH2PO4; 24 NaHCO3; 5 HEPES; 13 glucose; 2 MgSO4'; 2 CaCl2, oxygenated with carbogen gas (95% O2 and 5% CO2) to pH 7.3 - 7.4. To microdissect the PVN, slices were placed in a petri-dish containing ice-cold, oxygenated ACSF solution and the PVN was identified using the 3rd ventricle and other structural markers. Following dissection, tissue was placed into 2.6 mL equilibrated Papain DNase-I solution. The dissociation protocol was similar to Chevée et al., 2018, itself an adaptation from the trehalose-enhanced neuronal isolation protocol(*47*) using the Worthington Papain Dissociation System (Worthington Biochemical Corporation). The following modifications from Chevée et al., 2018 were made: a single low speed centrifugation (300xg for 5 min) was performed after dissociation, and the pellet was resuspended in 250 µL media (DMEM, 5% trehalose (w/v), 25 µM AP-V, 0.4 mM kynurenic acid, 6 µL of 40 U/µl RNase inhibitor) and 250 uL of EBSS#2 (EBSS, 25 mM AP-V, 100 mM

Kynurenic acid, ovomucoid protease inhibitor with BSA, DNase-I, 5% w/v Trehalose, 40 U/µl RNase inhibitor) at room temperature. Following resuspension, single-cell suspensions were introduced into a FACS machine (Beckman Coulter MoFlo Cell Sorter).

Neurons were sorted based on fluorescence (GFP+/FG- or GFP+/FG+) directly into individual wells of a 96-well plate containing 2 µL Smart-Seq2 lysis buffer + RNAase inhibitor, 1µL oligo-dT primer, and 1 µL dNTPs(*48*). Three plates were collected across 2 separate dates (Plate 1: n = 1 mouse, 32 GFP+/FG- neurons and 47 GFP+/FG+ neurons; Plate 2: n = 1 mouse, 32 GFP+/FG- neurons and 33 GFP+/FG+ neurons; Plate 3: n = 5 mice, 25 GFP+/FG- neurons and 48 GFP+/FG+ neurons). Following the sort, plates were briefly spun-down in a table-top microcentrifuge and immediately placed on dry ice. Single-cell lysates were kept at -80°C until cDNA library preparation. Library preparation and amplification of single-cell samples was performed using a modified version of the Smart-Seq2 protocol(*22*).

Individual libraries were quality controlled, pooled, and sequenced on two lanes of an Illumina HiSeq 2500 sequencer to an average depth of 1,424,499 ±77,731 paired-end 50bp reads per neuron. Reads were aligned to the reference mouse genome (mm10) using the Hisat 2 aligner and transcript abundances were estimated using CuffNorm (*49*) against the mouse GENCODE reference transcriptome (vM8).(*49*) Relative abundances were converted to absolute estimates of gene expression using the Monocle2 CENSUS approach and used as input for the dpFeature workflow (*49*). 146 neurons passed our quality assessment, and we identified a total of 10,172 genes with detectable expression in at least 5 neurons. For those cells that passed QC, the mean mRNA-copies per cell was 27,551 and mean number of genes detected across all cells was 3,713. A Uniform

Manifold Approximation and Projection (UMAP)(*50*) embedding was obtained by collapsing the top 50 principal components learned on the top 1000 genes with highest residuals to the Monocle model fit. All data are archived in the Gene Expression Omnibus (GEO Accession number GSE147092). Raw and processed scRNA-Seq data are also publicly available at https://github.com/gofflab/OT_neuron_study_2020

*In situ* hybridization chain reaction

Experiments were performed on 3 week-old FG injected male wild-type C57BL6/J mice obtained from Jackson Laboratories. Brains were extracted and immediately frozen in OCT with liquid nitrogen. Brains were sectioned on a cryostat at 7 µm. Serial sections were collected onto charged slides, briefly washed in PBS and then fixed in 4% PFA for 10 mins. Slides were then washed twice in PBS, transferred to 70% ethanol, and kept at 4˚C. *In situ* HCR v3.0 was performed using the protocols previously detailed (*24*). Probe hybridization and amplification steps were both performed over night (12-16 hours). *In situ* probes (**Table S2**) were design using publicly available software (*51*).

Immunohistochemistry

Prior to immunostaining, sections were mounted on slides and rinsed 4 x 10 min in PBS followed by 1 hr in blocking solution (0.5% Triton X-100, 10% horse serum, 0.2% bovine serum albumin in PBS). Antibodies were diluted in PBS containing the following: 0.5% Triton X-100, 1% horse serum, 0.2% bovine serum albumin. Primary antibodies were applied overnight at room temperature (RT), and after rinsing slides in PBS, secondary antibodies were applied for 2 hr at

RT. With the exception of FMRP immunohistochemistry, OT neurons were labeled using the anti-OT neurophysin antibody PS38 (gift of Dr. Harold Gainer; 1:150). PS38 was detected using Alexa 488, (magnocellular and parvocellular distribution; Life Technologies; donkey anti-mouse; 1:1000) or Alexa 647 (GFP or tdTomato colocalization; Life Technologies; goat-anti mouse or donkey-anti mouse respectively; 1:1000). FluoroGold was labeled with an anti-fluorogold antibody, (Fluorochrome; 1:100), and detected using Alexa 647 (Life Technologies; donkey anti-rabbit; 1:1000). GFP signal was amplified using the anti-GFP antibody, ab13970, (ABCAM; 1:1000) and visualized using Alexa 488 (Life Technologies; goat-anti chicken; 1:1000). tdTomato signal was amplified using the anti-mCherry antibody AB0040 (SICGEN; 1:4000) and visualized using Alexa 555 (Life Technologies; donkey-anti goat; 1:1000). For post-hoc identification of electrophysiologically recorded neurons, Neurobiotin was detected using streptavidin-conjugated Alexa 350 (Life Technologies; 1:800).

FMRP co-labeling experiments were conducted as described above with the following exceptions. After rinsing in PBS, slides were transferred to a hot (90°C) sodium citrate (10 mM Sodium Citrate, 0.05% Tween 20, pH 6.0) bath for antigen retrieval (30 min) followed by 1hr in blocking solution at RT. Sections were labeled with anti-FMRP 2F5-1 ((*52*) Developmental Studies Hybridoma Bank; 1:1) which was detected using Alexa 647 (Jackson ImmunoResearch; goat anti-mouse IgG2b; 1:500). OT neurons were labeled with the anti-oxytocin antibody VA10(*53*) (gift of Dr. Harold Gainer; 1:1000) and detected using Alexa 488 (Life Technologies; donkey anti-rabbit; 1:1000).

Histology

Following transcardial perfusion with 1M PBS and 10% formalin, brains were kept in formalin at 4˚C overnight and then transferred to PBS. To prepare them for re-sectioning, acute brain slices used for electrophysiology were placed in 10% formalin immediately after recording, and then transferred to PBS. For all experiments in which fluorescent colocalization was analyzed, serial brain sections (20 μm thickness) were cut using a cryostat following cryoprotection of tissue with a 30% sucrose solution containing 0.01% azide. Following immunostaining, images were acquired using an EVOS or Olympus BX41 microscope with 4x, 10x, and 40x objectives and analyzed using ImageJ.

Statistics and Data Analysis

All behavioral and electrophysiological analyses were performed with MATLAB (Mathworks) or Prism (GraphPad). Comparisons between experimental manipulations were made using a two-tailed, Students t-test (paired or unpaired, as appropriate), with $P < 0.05$ considered significant. For k-means clustering analysis, two clusters were specified, as this number of clusters was determined to be optimal by the elbow method. Electrophysiological feature values used for the clustering analysis were z-scored across all neurons, while the clustering algorithm was trained only on the values from neurons from Fig. 1.

All scRNAseq analysis was performed in R/Bioconductor following alignment with Hisat2. Log2 expression estimates (with a pseudocount of 1) of high-variance genes were used as input for PCA analysis and UMAP clustering of individual cells. After cluster assignment, differential expression testing was performed using the Monocle2 VGAM model comparison test(*23*) (FDR

0.1%, Monocle2 test, Benjamini-Hochberg corrected). Gene set enrichment was computed using the hypergeometric test unless stated otherwise. All code is available upon request.

**Fig. S1.**

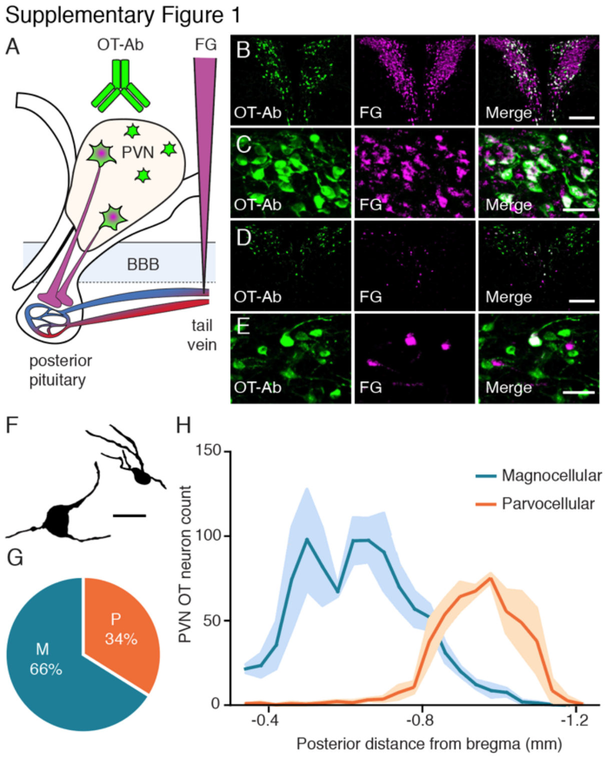


Distribution of OT neuronal subtypes across the rostral caudal axis of the PVN. **A,** Diagram illustrating labeling strategy. **B-E**, Example images of rostral (B,C) and caudal (D,E) sections of the PVN at low magnification (B,D, scale bar 200 μm) and high magnification (c,e, scale bar 50 μm) showing OT-Ab (green, left), FG (magenta, center) and merged (right) images. **F,** Drawing of a magnocellular (left) and parvocellular (right) OT neuron identified by i.v. FG labeling. Scale bar indicates 20 μm. **G,** Magnocellular OT neurons account for 65.9 ± 2.6% and parvocellular OT neurons account for 34.1 ± 2.6% of PVN OT neurons (n = 3 animals; 1439.3 ± 48.3 OT neurons per animal). **H,** Magnocellular OT neurons are predominantly located in the rostral PVN, while parvocellular OT neurons are predominantly located in the caudal PVN. Data represented as mean ± sem.

**Fig. S2.**

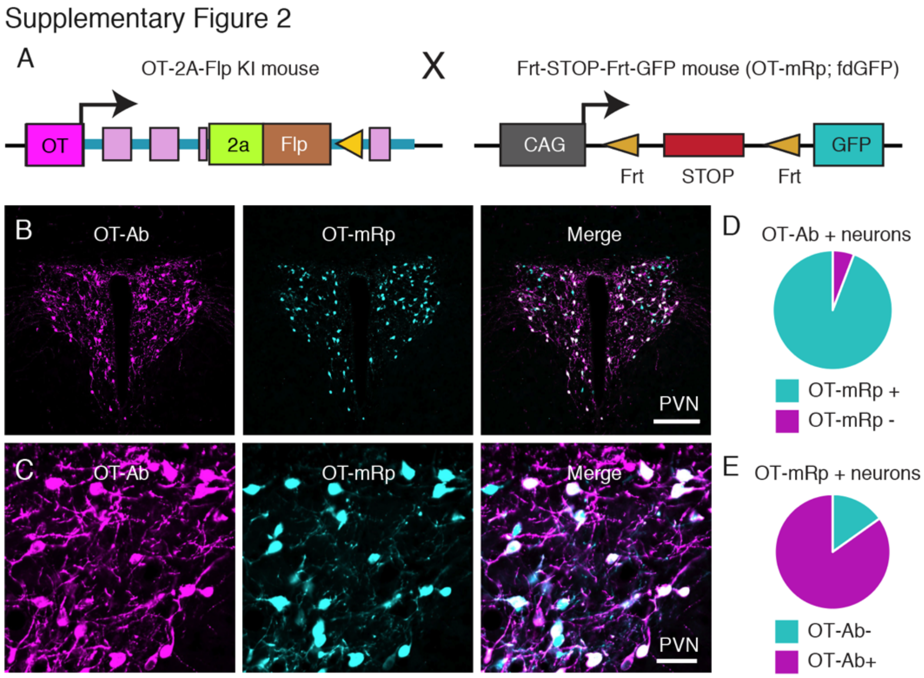


Validation of OT-2A-Flp::fdGFP cross to label OTergic neurons. **A,** Breeding strategy for OT-2A-Flp driver mice crossed to fdGFP reporter mice. **B**, OT antibody (OT-Ab, magenta left), GFP (OT-mRP, cyan, center), and merged (right) image of the PVN at low magnification (scale bar 200 µm). **C**, OT-Ab (magenta left), OT-mRP (cyan, center), and merged (right) image of the PVN at high magnification (scale bar 50 µm). **D,E,** Quantification of the efficacy of the mouse reporter strategy (n = 1 male mouse, P36, 862 neurons counted). OT-Ab positive neurons are 94.2% mRp positive (D). OT-mRp positive neurons are 84.9% OT-Ab positive (E).

**Fig. S3.**

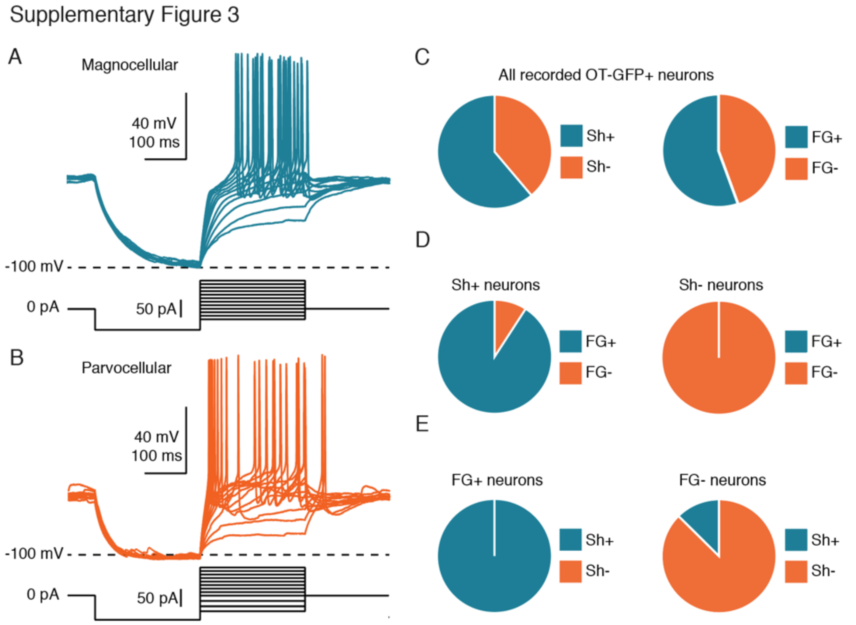


Electrophysiological protocol to distinguish magnocellular and parvocellular OT neurons and cross-validation with FG labeling. **A,** Overlay of 12 electrophysiological traces in which a magnocellular OT neuron was hyperpolarized to -100 mV, and depolarized from that membrane potential with current steps of increasing amplitude. **B,** Overlay of 12 electrophysiological traces in which a parvocellular OT neuron was hyperpolarized to -100 mV, and depolarized from that membrane potential using current steps of increasing amplitude. **C-E** The proportions of neurons identified as magnocellular or parvocellular electrophysiologically are consistent with those identified in **Fig. S1**. **C,** Left; 61% (n = 11 of 18) of sampled neurons were Sh+, and 39% (n = 7 of 18) were Sh-. Right; 56% (n = 10 of 18) sampled neurons were FG+, and 44% (n = 8 of 18) were FG-. **D,** Left; 91% (n = 10 of 11) Sh+ neurons were FG+. Right; 100% (n = 7 of 7) Sh- neurons were FG-. **E**, Left; 100% (n = 10 of 10) FG+ neurons were also Sh+. Right; 88% (n = 7 of 8) FG- neurons were Sh-.

**Fig. S4.**

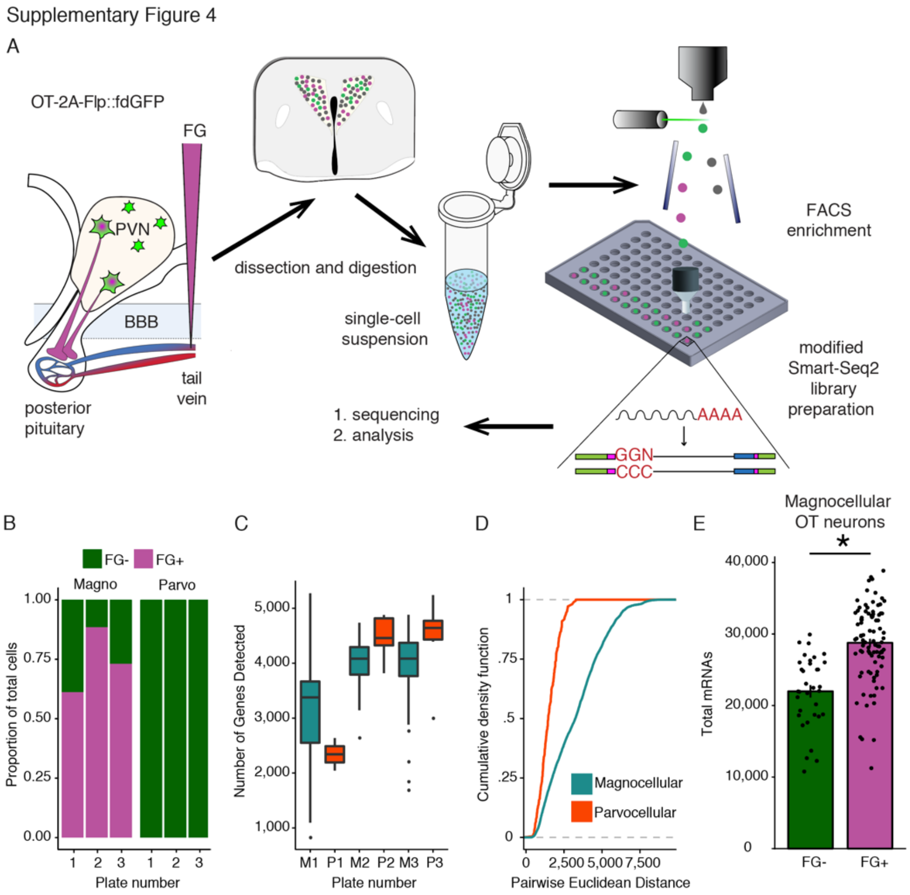


scRNASeq of oxytocin expressing neurons in the PVN. **A,** Schematic overview of experiment. **B,** Proportion of each cell type on each plate accounting for FG signal. **C,** Number of genes detected in each cell type by technical batch. **D,** Empirical Cumulative Distribution Functions for all expressed genes in the two cell types indicated greater intra-cell type heterogeneity within magnocellular neurons ($p = 1.665e^{-15}$). **E,** Magnocellular OT neurons without detectable i.v. FG labeling contain fewer total mRNAs than magnocellular OT neurons with detectable i.v. FG labeling ($t_{(123)} = 6.4904$ $p = 1.891^{e\text{-}09}$). * indicates $p < 0.05$; K-S test (D) or Pearson's product moment test (E) for statistical comparisons.

**Fig. S5.**

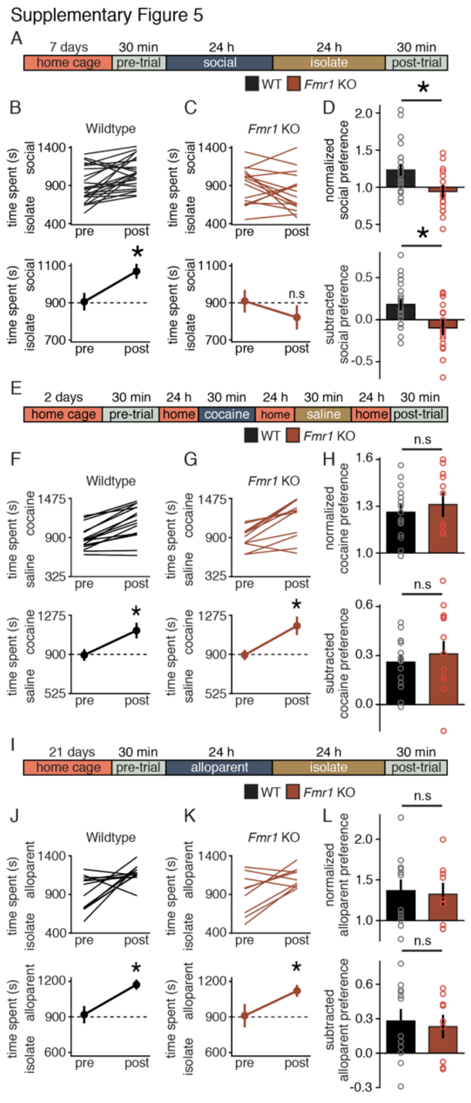


*Fmr1* KO mice exhibit social domain specific impairments in reward learning. **A,** Timeline for social CPP. **B,C,** Individual (top) and average (bottom) data for WT mice (B) and *Fmr1* KO mice (C). WT ($t_{(23)} = 3.586$; $p = 0.002$) but not *Fmr1* KO mice ($t_{(15)} = 1.324$; $p = 0.205$) exhibit social CPP. **D,** Normalized ($t_{(38)} = 2.917$; $p = 0.006$) and subtracted ($t_{(38)} = 3.23$; $p = 0.003$) comparisons of social preference reveal a difference in the magnitude of social CPP between WT and *Fmr1* KO mice. **E,** Protocol and timeline for cocaine CPP. **F,G** Individual (top) and average (bottom) data for WT mice (f) and *Fmr1* KO mice (g). Both WT ($t_{(14)} = 6.454$; $p < 0.0001$) and *Fmr1* KO mice

($t_{(10)} = 4.297$; $p = 0.002$) exhibit cocaine CPP. **H,** Normalized ($t_{(24)} = 0.609$; $p = 0.548$) and subtracted ($t_{(24)} = 0.644$; $p = 0.525$) comparisons of cocaine preference reveal a similar magnitude of cocaine CPP in WT and *Fmr1* KO mice. **I,** Protocol and timeline for alloparent CPP. **J,K** Individual (top) and average (bottom) data for WT mice (F) and *Fmr1* KO mice (G). Both WT ($t_{(11)} = 2.931$; $p = 0.014$) and *Fmr1* KO mice ($t_{(8)} = 2.488$; $p = 0.038$) exhibit alloparent CPP. **L,** Normalized ($t_{(19)} = 0.234$; $p = 0.817$) and subtracted ($t_{(19)} = 0.354$; $p = 0.727$) comparisons of alloparent preference reveal a similar magnitude of alloparent CPP in WT and *Fmr1* KO mice. Data represented as mean ± sem. * indicates $p < 0.05$; Student's t-test (two-tailed; paired (B,C,F,G,J,K) or unpaired (D,H,L)) for statistical comparisons.

**Fig. S6.**

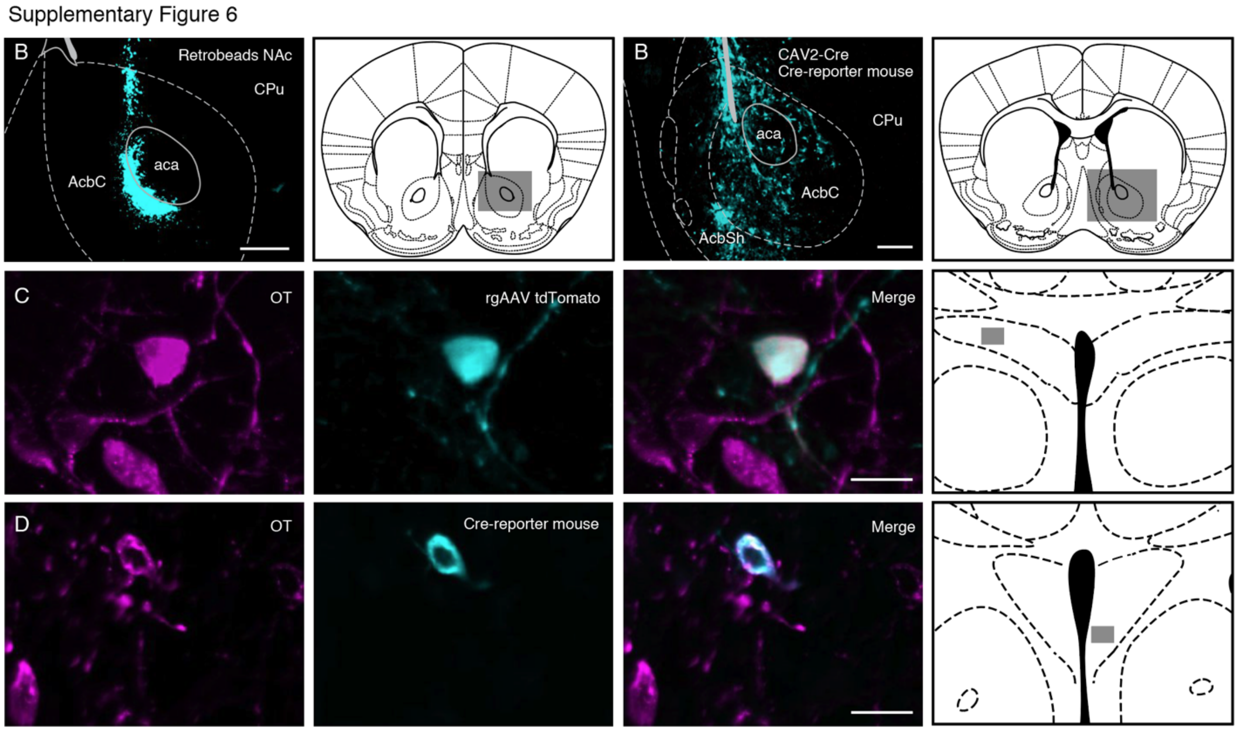


Injection targeting and viral expression. **A,** Retrobead injection targeted to NAc (left) and atlas image depicting the area captured in the image (right). **B,** CAV2-Cre infection induces Cre-mediated recombination. (left) CAV2-Cre injection targeted to NAc of a tdTomato Cre-reporter mouse and (right) atlas image depicting the area captured in the image. **C,** retro-AAV infects PVN OT neurons that innervate NAc. From left to right: OT-Ab; retro-AAV-tdTomato expression; Merge; atlas image indicating the location captured in images to the left. **D,** retro-AAV-fdCre can induce Flp-dependent Cre-mediated recombination in PVN OT neurons. From left to right: OT-Ab; Cre-mediated tdTomato expression; Merge; atlas image indicating the location captured in images to the left. Atlas images adapted from Paxinos and Franklin, 2001. Scale bars indicate 200 µm in low magnification images, and 20 µm in high magnification images.

**Fig. S7.**

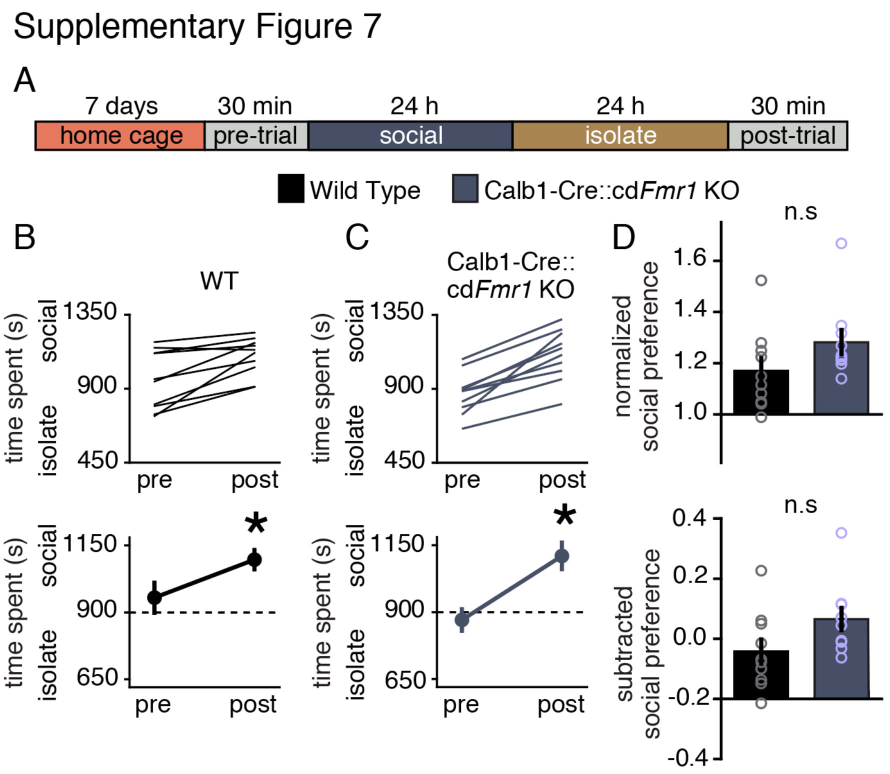


*Fmr1* knock down in magnocellular OT neurons does not impair social reward learning. **A,** Protocol and timeline for social CPP. **B,C,** Individual (top) and average (bottom) data for WT (B) and *Calb1*-IRES-Cre::cd*Fmr1* KO mice (C). WT ($t_{(9)}$ = 3.901; p = 0.004) and *Calb1*-IRES-Cre::cd*Fmr1* KO mice ($t_{(9)}$ = 7.146; p > 0.0001) mice exhibit social CPP. **D,** Normalized ($t_{(18)}$ = 1.642; p = 0.118) and subtracted ($t_{(18)}$ = 1.965; p = 0.065) comparisons of social preference reveal that WT and *Calb1*-IRES-Cre::cd*Fmr1* KO mice exhibit similar magnitude social CPP. Data represented as mean ± sem. * indicates p < 0.05; Student's t-test (two-tailed; paired (B,C) two-tailed; unpaired(D)) for statistical comparisons.

**Fig. S8.**

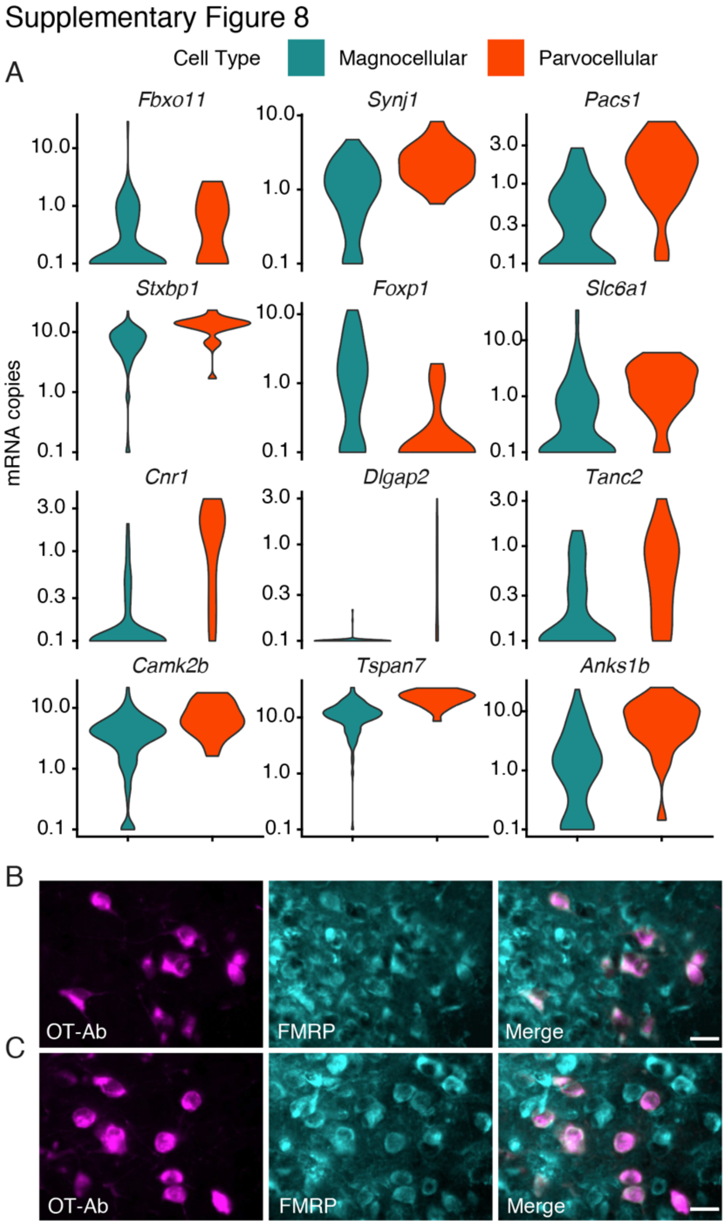


Differentially expressed ASD risk genes that are binding targets of FMRP. **A,** Violin plots for the set of 12 FMRP binding targets that are also ASD risk genes. Of the 12 genes, 10 are enriched in parvocellular OT neurons. **C,D,** FMRP is expressed in OT neurons in the rostral (C) and caudal (D) PVN. Scale bar 20 µm. * indicates $p < 0.05$; Monocle Likelihood Ratio Test for statistical comparisons.

**Fig. S9.**

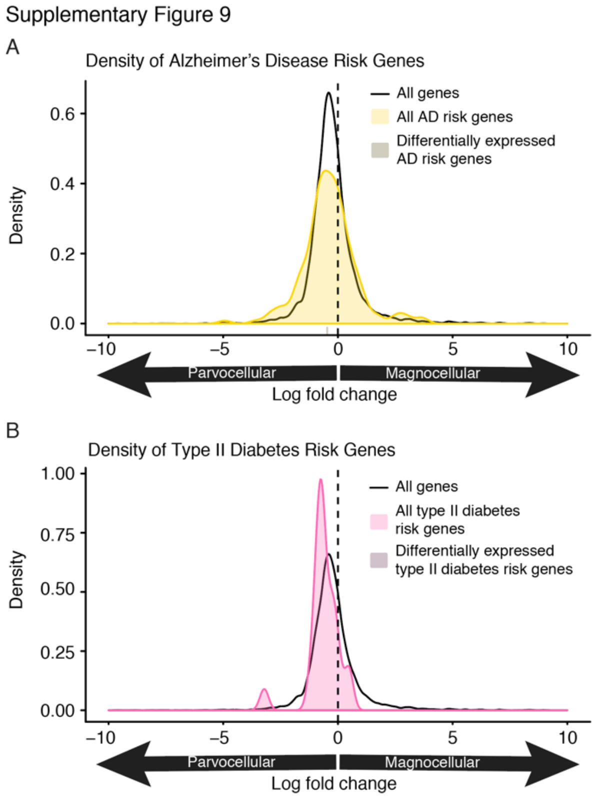


Alzheimer's disease and Type II Diabetes risk genes are not differentially expressed in magnocellular and parvocellular OT neurons. **A,** Density of the log2 fold change for (black) all expressed genes, (light yellow) all Alzheimer's disease risk genes, and (dark yellow) all significantly differentially expressed (FDR 0.1%) Alzheimer's disease risk genes. Alzheimer's disease risk genes are not enriched in magnocellular or parvocellular OT neurons ($p = 0.99$). **B,** Density of the log2 fold change for (black) all expressed genes, (light pink) all type II diabetes risk genes, and (dark magenta) all significantly differentially expressed (FDR 0.1%) type II diabetes risk genes. Type II diabetes risk genes are not enriched in magnocellular or parvocellular OT neurons ($p = 0.33$). Vertical tick mark below the AD density plots indicates the single differentially expressed AD risk gene (A). Note that lack of differentially expressed type II diabetes risk genes (B).

**Fig. S10.**

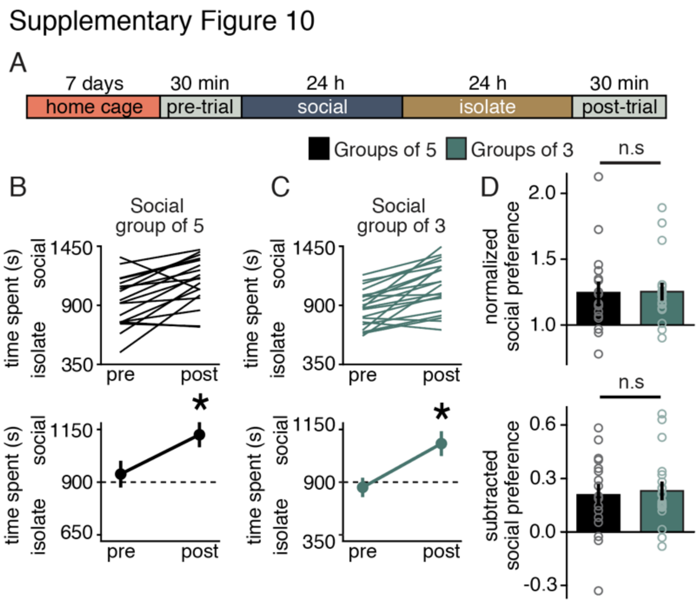


Social groups of 5 or 3 mice exhibit similar magnitude social CPP**. A,** Protocol and timeline for social CPP. **B,C,** Individual (top) and average (bottom) data for mice housed and conditioned in groups of 5 (B) and groups of 3 (C). Groups of 5 ($t_{(17)} = 3.988$; $p = 0.001$) and groups of 3 ($t_{(19)} = 5.287$; $p > 0.0001$) mice exhibit social CPP. **D,** Normalized ($t_{(36)} = 0.079$; $p = 0.937$) and subtracted ($t_{(36)} = 0.341$; $p = 0.735$) comparisons of social preference reveal that mice conditioned in groups of 5 or groups of 3 exhibit similar magnitude social CPP. Data represented as mean ± sem. * indicates $p < 0.05$; Student's t-test (two-tailed; paired (B,c) two-tailed; unpaired(D)) for statistical comparisons.

**Table S1.**
Differentially expressed genes. (Excel)

**Table S2.**

| Probe Name | Probe Sequence |
|---|---|
| Calbindin_CDS_HCR_B1_P1_odd | GAGGAGGGCAGCAAACGGAATCCTTCCAGGTAACCACTTC |
| Calbindin_CDS_HCR_B1_P2_odd | GAGGAGGGCAGCAAACGGAATCTGTCCATATTGATCCACA |
| Calbindin_CDS_HCR_B1_P3_odd | GAGGAGGGCAGCAAACGGAAGCTGTGGTCAGTATCATACT |
| Calbindin_CDS_HCR_B1_P4_odd | GAGGAGGGCAGCAAACGGAATTCCCACACATTTTGATTCC |
| Calbindin_CDS_HCR_B1_P5_odd | GAGGAGGGCAGCAAACGGAAATCTATGTATCCGTTGCCAT |
| Calbindin_CDS_HCR_B1_P6_odd | GAGGAGGGCAGCAAACGGAAAGCTTCCCTCCATCCGACAA |
| Calbindin_CDS_HCR_B1_P1_even | GGATCAAGTTCTGCAGCTCCTAGAAGAGTCTTCCTTTACG |
| Calbindin_CDS_HCR_B1_P2_even | ATTCCTATTTTTCCATCATCTAGAAGAGTCTTCCTTTACG |
| Calbindin_CDS_HCR_B1_P3_even | GTTCCTCGGTTTCGATGAAGTAGAAGAGTCTTCCTTTACG |
| Calbindin_CDS_HCR_B1_P4_even | CTCAAAAGCCTTATTGAACTTAGAAGAGTCTTCCTTTACG |
| Calbindin_CDS_HCR_B1_P5_even | GCAAAGCATCCAGCTCATTTTAGAAGAGTCTTCCTTTACG |
| Calbindin_CDS_HCR_B1_P6_even | AAGAGCAAGGTCTGTTCGGTTAGAAGAGTCTTCCTTTACG |
| OXT_CDS_HCR_B3_P1_odd | GTCCCTGCCTCTATATCTTTAGCAAGCGAGACTGGGGCAG |
| OXT_CDS_HCR_B3_P2_odd | GTCCCTGCCTCTATATCTTTTCTGGATGTAGCAGGCCGAG |
| OXT_CDS_HCR_B3_P3_odd | GTCCCTGCCTCTATATCTTTGTCCGAAGCAGCGTCCTTTG |
| OXT_CDS_HCR_B3_P4_odd | GTCCCTGCCTCTATATCTTTGAAGGCAGGTAGTTCTCCTC |
| OXT_CDS_HCR_B3_P5_odd | GTCCCTGCCTCTATATCTTTAGGCGGGGTCTGTGCGGCAG |
| OXT_CDS_HCR_B3_P1_even | AGAGCCAGTAAGCCAAGCAGTTCCACTCAACTTTAACCCG |
| OXT_CDS_HCR_B3_P2_even | CTCTTGCCGCCCAGGGGGCATTCCACTCAACTTTAACCCG |
| OXT_CDS_HCR_B3_P3_even | TCGTCCGCGCAGCAGATGCTTTCCACTCAACTTTAACCCG |
| OXT_CDS_HCR_B3_P4_even | CTTCTGGCCAGACTGGCAGGTTCCACTCAACTTTAACCCG |
| OXT_CDS_HCR_B3_P5_even | GAGAAGGCAGACTCAGGGTCTTCCACTCAACTTTAACCCG |
| Cnr1_CDS_HCR_B1_P1_odd | GAGGAGGGCAGCAAACGGAACAAGGCCGTCTAAGATCGACTTCAT |
| Cnr1_CDS_HCR_B1_P2_odd | GAGGAGGGCAGCAAACGGAAATCTTCGTACTGAATGTCATTTGAG |
| Cnr1_CDS_HCR_B1_P3_odd | GAGGAGGGCAGCAAACGGAACATAAAATTCTCCCCACACTGGATG |
| Cnr1_CDS_HCR_B1_P4_odd | GAGGAGGGCAGCAAACGGAAGGAGACTGCGGGAGTGAAGGATGAC |
| Cnr1_CDS_HCR_B1_P5_odd | GAGGAGGGCAGCAAACGGAAGTGGAACACGTGGAAGTCAACAAAG |
| Cnr1_CDS_HCR_B1_P6_odd | GAGGAGGGCAGCAAACGGAACCACATCAAGCAAAAGGCCACTACG |
| Cnr1_CDS_HCR_B1_P7_odd | GAGGAGGGCAGCAAACGGAAACAGATTGCAGCTTCTTGCAGTTCC |
| Cnr1_CDS_HCR_B1_P8_odd | GAGGAGGGCAGCAAACGGAACCGATCCAGAACATCAGGTAGGTTT |
| Cnr1_CDS_HCR_B1_P9_odd | GAGGAGGGCAGCAAACGGAACAGAGGGCCCCAGCAGATGATCAAC |
| Cnr1_CDS_HCR_B1_P10_odd | GAGGAGGGCAGCAAACGGAACGTCTTGATAAGCTTGTTCATCTTC |
| Cnr1_CDS_HCR_B1_P11_odd | GAGGAGGGCAGCAAACGGAACATGAAGGGAACATGCTGCGGAAAG |
| Cnr1_CDS_HCR_B1_P1_even | TGGTGATGGTACGGAAGGTGGTATCTAGAAGAGTCTTCCTTTACG |
| Cnr1_CDS_HCR_B1_P2_even | TAATTTGGATGCCATGTCTCCTTTGTAGAAGAGTCTTCCTTTACG |
| Cnr1_CDS_HCR_B1_P3_even | ATTCAGAATCATGAAGCACTCCATGTAGAAGAGTCTTCCTTTACG |
| Cnr1_CDS_HCR_B1_P4_even | CAATGAAGTGGTAGGAAGGCCTGCATAGAAGAGTCTTCCTTTACG |
| Cnr1_CDS_HCR_B1_P5_even | CAGAAACACATTGGGACTATCTTTGTAGAAGAGTCTTCCTTTACG |
| Cnr1_CDS_HCR_B1_P6_even | CAACACAGCAATTACTATTGCAATATAGAAGAGTCTTCCTTTACG |

| | |
|---|---|
| Cnr1_CDS_HCR_B1_P7_even | TCAATGAGTGGGAAGATGTCTGAGCTAGAAGAGTCTTCCTTTACG |
| Cnr1_CDS_HCR_B1_P8_even | ATGAACAGCAACAGCACACTGGTGATAGAAGAGTCTTCCTTTACG |
| Cnr1_CDS_HCR_B1_P9_even | AAAGACATCATACACCATGATCGCATAGAAGAGTCTTCCTTTACG |
| Cnr1_CDS_HCR_B1_P10_even | GCAGAGCATACTACAGAAGGCAAACTAGAAGAGTCTTCCTTTACG |
| Cnr1_CDS_HCR_B1_P11_even | TTATCTAGAGGCTGCGCAGTGCCTTTAGAAGAGTCTTCCTTTACG |

Hybridization chain reaction version 3.0 (HCR 3.0) probe design.

**Table S3.**
Differentially expressed ASD risk genes. (Excel)

**Table S4.**
Differentially expressed FMRP binding partners. (Excel)
